# Decoding the causal drivers of spatial cellular topology

**DOI:** 10.1101/2025.03.19.644241

**Authors:** Prannav Shankar, Huan Liang, Uthsav Chitra, Rohit Singh

## Abstract

Decoding how cells influence and communicate with each other in space is fundamental for understanding tissue organization. However, existing approaches either overlook spatial context entirely or rely solely on local cell-cell adjacency, failing to capture how global tissue topology shapes cellular communication. Here, we present GLACIER, which introduces *spatial Granger causality* to infer transcriptional and signaling relationships that emerge from tissue organization. By combining GASTON’s global isodepth coordinate with Velorama’s graph-based causal inference framework, we enable bidirectional inference of regulatory relationships along spatial axes, identifying transcription factor-target interactions and ligand-receptor pairs that operate across spatial domains. Applying GLACIER to single-cell spatial transcriptomics data from the mouse cerebellum, we identify both continuous within-cell-type regulatory gradients and discontinuous drivers at layer interfaces, while distinguishing between forward and backward cellular communication along the isodepth axis. Our approach reveals how tissue architecture directs patterns of cellular communication, providing a framework for understanding spatially-encoded regulatory programs.

**Software availability:** https://github.com/rohitsinghlab/glacier.

## 1 Introduction

Multicellular organisms depend on precisely orchestrated cell-cell interactions that unfold across multiple spatial scales, including both direct, short-range interactions and indirect, longer-range communication. For example, cells directly communicate with one another by releasing ligand molecules that bind to receptors on neighboring cells, triggering immediate signaling changes. More broadly, transcription factors (TFs) can initiate regulatory programs that cascade across cellular neighborhoods and lead to coordinated changes in gene expression programs in larger domains of a tissue [1]. Understanding these spatially-encoded cellular interactions is crucial for decoding many fundamental biological processes including tissue development [2], immune response [3], wound healing [4], and cancer progression [5].

Recent advances in spatial transcriptomics (ST) offer unprecedented potential for characterizing cellular interactions in tissues. Unlike traditional single-cell RNA sequencing (scRNA-seq) which dissociates cells before measurement and loses spatial information, ST technologies measure both gene expression *and* the spatial location of individual cells in tissue slices. However, there are two key challenges in the inference of cellular interactions from current ST datasets. First, transcript counts in ST data are highly sparse, making it difficult to reliably estimate short-range, *local* correlations in gene expression [6, 7]. For example, whole-transcriptome technologies such as Slide-SeqV2 [8] and Stereo-seq [9] measure a median of less than 500 unique molecular identifiers (UMIs) per cell. Second, and more fundamentally, understanding how cells communicate over long scales requires knowledge of the *global* geometry of the tissue and the spatial axis along which regulatory signals propagate.

Current computational methods for inferring cellular or regulatory interactions from ST data do not leverage global tissue organization. Nearly all methods implicitly assume a uniform interaction likelihood between neighboring cells by representing these local relationships through cell-cell adjacency graphs, and they look for correlated TF-gene or ligand-receptor expression along edges of such an adjacency graph [6, 10, 11, 12, 13, 7]. However, this oversimplified view fails to capture how intercellular signals propagate in specific directions determined by tissue architecture. For instance, endothelial cells communicate along paths defined by tissue cytoskeleton [14], while the Notch signaling pathway operates bidirectionally along particular spatial axes during development [15]. Accurate inference of cellular interactions thus requires incorporating global tissue structure to understand the directionality of gene expression variation.

GASTON [16] addresses this need by learning a “topographic map” of tissue from ST data. This map is defined by a spatial coordinate called “isodepth” which varies globally across the tissue slice, and measures the direction of maximum variation in gene expression. It allows inference of directional cellular interactions by structuring the tissue into a hierarchical framework. This approach helps distinguish between local interactions and broader regulatory signals dictated by global tissue topology. In the mouse cerebellum, GASTON successfully revealed both discrete cellular domains (layers) and continuous gradients of gene activity, providing the first comprehensive map of spatial gene expression in this tissue. However, GASTON models each the variation of gene independently across the isodepth coordinate and does not account for gene-gene interactions, leaving open the challenge of inferring cellular communication and regulatory relationships from ST data.

We introduce *spatial Granger causality*, a fundamentally new approach for learning cellular interactions from ST data. The key insight is that tissue organization, as captured by GASTON’s isodepth, creates natural directions of information flow that can be represented as a directed acyclic graph (DAG). While classical Granger causality required data to follow a linear time series, recent advances [17, 18, 19] have enabled Granger causal inference on partial orderings represented by DAGs. Building on this foundation, we present GLACIER (Granger-Led Analysis of Cellular Isodepth and Expression Regulation), which combines GASTON’s global spatial coordinate with Velorama’s DAG-based nonlinear Granger causality to identify TF-gene and ligand-receptor relationships that propagate along spatial axes. GLACIER systematically captures how regulatory information flows through tissue structure, enabling directional inference of cellular signaling interactions. Applied to Slide-SeqV2 data of mouse cerebellum [8, 20], GLACIER identifies directional transcriptional programs within layers, revealing that oligodendrocytes predominantly regulate neuronal connectivity in one direction. At the boundary between the granular and Purkinje-Bergmann layers, GLACIER reports regulatory activity that is distinct from what is seen in either layer individually, suggesting that inter-layer communication operates through specialized mechanisms rather than a simple gradient. These findings demonstrate how integrating spatial organization with causal inference can reveal fundamental principles of tissue architecture and cellular communication.

## 2 Methods

### 2.1 Granger causality in time-series

Granger causality is a statistical framework for inferring causal relationships in time-series data by assessing whether past values of one variable x (predictor) improve the prediction of another variable y (target) beyond what can be explained by the latter’s own past values. The key principle is that a cause must precede its effect, and predictive relationships between variables may therefore indicate potential causality. Granger causal interactions are statistical estimates, best interpreted as causal hypotheses to be prioritized for experimental validation; for example, a Granger causal relationship between x and y may be indirect, mediated through a latent and potentially unobserved variable.

### Linear Granger causality

Given two time series of length *N*, 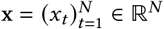 and 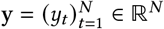,we say that they follow a *vector autoregressive (VAR)* model with *L >* 0 lags if

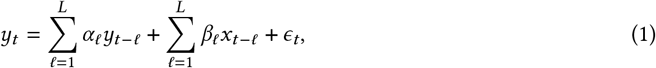

where *ϵ*_*t*_ is a noise term, and *α*_*𝓁*_, *β*_*𝓁*_ ℝ are coefficients that describe the contribution of the observations *y*_*t*− *𝓁*_, *x*_*t* −*𝓁*_, respectively, at the *𝓁*-th *lag*, i.e. the (*t* − 𝓁)-th time. The value *L >* 0 indicates how far back in time past observations influence the observation *y*_*t*_, i.e. the *maximum lag*. VAR models are widely used for time-series analysis in finance and economics [21].

We say that x linearly *Granger causes* y if *β*_*𝓁*_ ≠ 0 for some lag 0 < *𝓁*≤ *L* [22]. The standard approach for determining linear Granger causality is through hypothesis testing, where the null hypothesis is *H*_0_ : *β*_*𝓁*_ = 0 for all lags *𝓁* = 1, …, *L*, e.g. using an F-test or likelihood ratio test [23, 24].

### Non-linear Granger causality

In many scenarios, the time series x, y are not linearly related. Thus, we generalize (1) and say that x and y follow a *non-linear vector autoregressive (NVAR)* model with *L* lags if

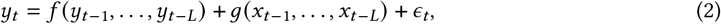

where *f, g* : ℝ^*L*^ → ℝ are functions— potentially parameterized by neural networks— that specify how the observation *y*_*t*_ at time *t* depends on the previous *L* observations *y*_*t* −1_, …, *y*_*t* −*L*_ and *x*_*t* −1_, …, *x*_*t* −*L*_, respectively, and *ϵ*_*t*_ is a noise term.

In a non-linear setting, Granger causality can be detected through two approaches: *ablation-* or *invariance-* based [17]. The approporiate choice depends on the problem setting and the kind of interactions modeled between predictor variables:

- **Ablation-based approach**. Causality is assessed by training two models: a full model ℳ_full_ that includes past values of both x and y, and a restricted model ℳ_reduced_ that omits x. If removing x significantly reduces predictive accuracy, x is deemed Granger-causal of y.
- **Invariance-based approach**. Instead of explicitly comparing models, this approach applies regularization to x-dependent terms. If removing x does not alter the model’s predictions (e.g., its coefficients shrink to zero under sparsity constraints), then x does not Granger-cause y. This method is computationally efficient and allows joint learning of multiple interactions. We note that applying it in a neural network setting requires special architecture and training considerations to encode the required sparsity constraints [17].

### 2.2 Spatial Granger causality

Classical Granger causality approaches require a total ordering on the *N* observations, and are not immediately applicable to *spatially structured* data where there is no obvious total ordering of locations in space.

Here we leverage recent extensions of Granger causality to *partially ordered* data described by a directed acyclic graph (DAG) [17, 18, 19]. To extend Granger causality to spatially structured data, we combine the DAG-structured Granger causality framework with a novel construction of a *spatial DAG*.

We start by describing the DAG-structured Granger causality framework of GrID-Net and Velorama [17, 18, 19]. Given a DAG *G* = *V, E* and observations x = (*x*_*v*_)_∈*v V*_ ∈ℝ^|*V*|^, y = (*y*_*v*_)_*v*∈ *V*_ ∈ℝ^|*V*|^ defined on each vertex *v ∈ V*, we say that the observations x, y follow a *DAG-structured vector autoregressive (DVAR)* model with *L* lags, modifying (2) as follows:

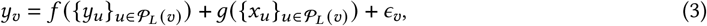

where P_*L*_ (*v*) is the set of vertices *u* ∈ *V* that are ancestors of *v* and are at most distance *L* from *v*. That is,

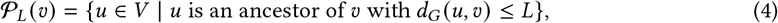

where *d*_*G*_ (*u, v*) is the length of the shortest directed path from *u* to *v* in *G*; and *f, g* : 2^ℝ^ →ℝ are functions that map each subset of the observations y, x, respectively, to a real number. As mentioned before, one can then infer causal relationships using an ablation or invariance-based approach. To estimate Granger causal relationships between individual genes and ATAC-seq peaks, GrID-Net used an ablation based approach [18, 25]. In contrast, when seeking to assess and select from multiple TFs (i.e., predictors) acting combinatorially, Velorama used an invariance-based approach with per-TF sparsity constraints [17].

#### DAG construction

We construct a spatial DAG using a *spatial potential* function, similar to how [17] constructs a DAG from a pseudotime function. Briefly, suppose we are given *N* spatial locations s_1_ = (*x*_1_, *y*_1_), …, s_*N*_ = (*x*_*N*_, *y*_*N*_) and a real number *d*_*i*_ ∈ℝ^*i*^ defined on each spatial location s_*i*_, which we call a *spatial potential*. We form a DAG *G* = (*V, E*) whose vertices *V* = {*s*_1_, …, *s*_*N*_} are the spatial locations, and whose edges *E* are those in the *k*-nearest-neighbors (*k*-NN) graph. Each edge (*i, j*) is oriented in the direction of increasing spatial coordinate; that is for any two arbitrary cells *i, j*, we include a directed edge *i* → *j* if *d*_*i*_ *> d* _*j*_. Such a construction guarantees that *G* is a DAG [17].

#### Isodepth

While our approach is generalizable to any spatial potential *d*_*i*_, in practice we use the spatial potential *d*_*i*_ learned by GASTON [16]. The GASTON spatial potential *d* is called the *isodepth* and is a 1-D latent coordinate describing the direction of maximum spatial variation in gene expression. For example, in layered tissues (e.g. the cerebellum tissue slice that we analyze in Section 3 and Figures 2,4), the isodepth *d* corresponds to the *layer depth*. See [16] for more details. We note that isodepth is just an example spatial potential and our method can be applied directly to perform causal analysis in the setting of other spatial potentials (e.g., radial distance from a blood vessel, germinal center, or tumor mass).

#### Inference

Since our inference objectives (i.e. identifying combinations of genes as regulators) are similar to Velorama, we follow its invariance-based approach to extend Granger causality to spatial transcriptomics. For each candidate regulator x = (*x*_*v*_) _*v*∈*V*_ (i.e. gene, out of *N*_*G*_ total candidates) and vertex *v* ∈*V* (i.e. cell) in *G*, we accumulate its expression across the 𝒫_*L*_ *v* ancestors of *v*. Together, these form the input to a neural network that predicts the expression of a target gene y = (*y*_*v*_) _*v* ∈*V*_. To enforce sparsity in the selection of regulators, we incorporate a regularization on the first layer of the neural network (Figure 1B,right). This acts as a feature selection mechanism, ensuring that only a subset of candidate regulators contribute to predicting the target gene. Specifically, the first hidden layer takes the form

**Figure 1:**
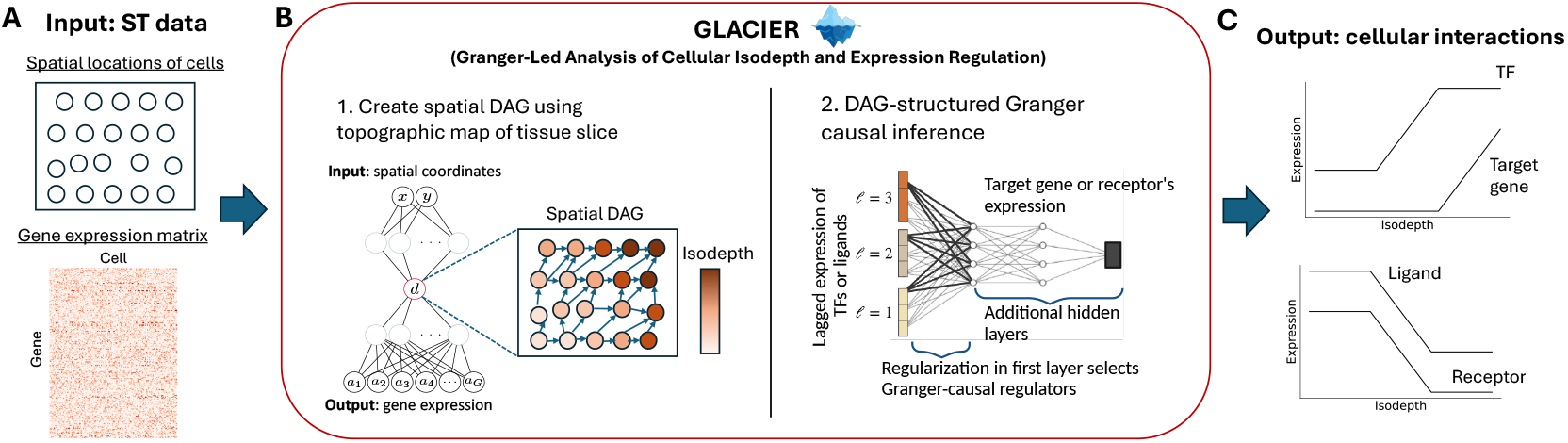
**(A)** The input to GLACIER is spatial transcriptomics (ST) data from a 2-D tissue slice, consisting of the spatial locations of measured cells and the gene expression matrix. **(B)** GLACIER (Granger-Led Analysis of Cellular Isodepth and Expression Regulation) first learns a topographic map of the tissue slice defined by an isodepth coordinate, and uses the topographic map to form a spatial DAG. GLACIER then performs directed acyclic graph (DAG)-structured Granger causal inference. **(C)** The output of GLACIER are cellular interactions: transcription factors (TFs) that are Granger causal for target genes, and ligand genes that are Granger causal for receptor genes.

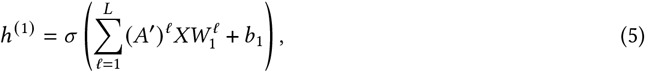

where (*A*′)^*𝓁*^ is the *𝓁*-th power of the adjacency matrix of 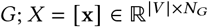 is the regulator expression matrix, where each column is a candidate regulator 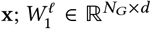 are trainable weight matrices with a lasso penalty; *d* is the dimensionality of the hidden layers; and *σ*(·) is a non-linear activation function.

Following [17], we apply a hierarchical regularization scheme that penalizes *W* ^*𝓁*^. Longer lags are more heavily penalized than shorter ones. This encourages robustness in selecting lagged interactions while reducing sensitivity to the choice of the maximum lag *L* [26]. To ensure sparsity, the model is trained using proximal gradient descent, which is well-suited for enforcing sparsity constraints. The regularization weight is specified by the hyperparameter *λ*. To select an appropriate *λ*, we perform a grid search over a range of values and ensemble the results across valid *λ* settings where some, but not all, weights remain nonzero. This enables GLACIER to infer spatially structured regulatory relationships that capture both local and long-range dependencies dictated by tissue architecture. We say that the *lag* of an interaction is the maximum number of non-zero weights. See [17] for details.

### 2.3 GLACIER algorithm and implementation

We present GLACIER, an algorithm for identifying Granger causal interactions from single-cell, spatial transcriptomics (ST) data (Figure 1A).

#### Input

The input to GASTON is ST data (A, S) which consists of

- an *N* × *G* gene expression matrix A = [*a*_*i,g*_] ∈ ℝ^*N* ×*G*^, where *a*_*i,g*_ is the expression of gene *g* ∈ {1, …, *G*} in spot/cell *i* ∈ {1, …, *N*},
- an *N* × 2 spatial location matrix S = [s_*i*_] ∈ ℝ^*N* ×2^, where s_*i*_ = (*x*_*i*_, *y*_*i*_) is the 2-D coordinate of spot/cell *i* for *i* = 1, …, *N*,

and a list ℒ ⊆ G ×*G* of putatively interacting genes; e.g. from a database of experimentally observed ligand-receptor interactions such as CellChatDB [27].

The ST data (A, S) is the standard output from applying a spatial sequencing or imaging technology to a 2-D tissue slice. Such spatial technologies include 10x Genomics Visium [28], Slide-Seq/Slide-SeqV2/Slide-tags [29, 8, 30], and MERFISH [31]. Different spatial technologies differ in the numbers *N, G* of spots and genes, respectively, that are measured, as well as their spatial resolution. See [32, 33, 34] for details. We also note that the list ℒ of putatively interacting genes is incomplete, i.e. not all interacting ligands and receptors are included in ℒ, and may also be biased towards genes that are well-studied (i.e. ascertainment bias).

#### Steps of GLACIER

GLACIER consists of the following steps. First, GLACIER uses GASTON [16] to compute the isodepth *d*_*i*_ for each cell *i* = 1, …, *N* and forms a spatial DAG *G*, as described in Section 2.2 (Figure 1B, left). For each pair (*g, h*) ∈ ℒ of candidate interacting genes, GLACIER uses the spatial DAG to test and report whether the observed gene-*g* expression values 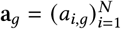 *Granger causes* the observed gene-*h* expression values 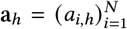 (Figure 1B, right), and reports all pairs (*g, h*) where this Granger causality relationship holds (Figure 1C).

We test for Granger causality using a version of Velorama [17] adapted to spatial data. We construct the spatial DAG using a *k*-NN graph (with *k* = 7) on each cell’s spatial coordinates, oriented using isodepth (Section 2.2). Notably, isodepth has the unique property that it can be *inverted* by redefining each cell’s isodepth as

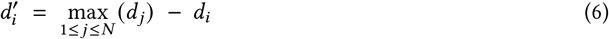

which reverses the ordering of isodepth among cells, allowing for analysis of information flow in the opposite direction.

#### Hyperparameter selection

Due to the sparsity of transcript counts from current ST technologies, several hyperparameter adjustments are necessary to ensure Velorama is able to run smoothly. We adjusted key hyperparameters as follows:

- **Learning Rate**: A higher learning rate than the default used in Velorama was sometimes necessary to achieve stable convergence. To balance this higher learning rate, we also implemented a learning rate scheduler.
- **Number of Epochs**: We also needed to increase the number of training epochs to allow the model sufficient time to reach an optimal solution due to the number of genes and learning rate scheduling.
- **Regularization (***λ*): Due to the sparsity of spatial transcriptomics data, we needed to sample from a broader range of *λ* values to ensure valid settings were obtained. As defined in [17], valid settings are those where between 5% and 95% of regularization weights remain nonzero.

To validate the robustness of our hyperparameter choices, we conducted a bootstrap-style analysis in which the dataset was randomly partitioned into three subsets. We selected hyperparameters that led to robust inference results across all three runs; see Appendix A for more details. We provide guidance on hyperparameter selection with the source code release. Running GLACIER on the full mouse cerebellum dataset, which consists of 9,985 cells, typically required *≈* 400 minutes on a single Nvidia A6000 GPU.

## 3 Results

We used GLACIER to identify cellular interactions in the mouse cerebellum using spatial transcriptomics (ST) data measured using the Slide-SeqV2 technology [8, 20]. The expression of 23, 096 transcripts was measured at 9, 985 spatial locations. The mouse cerebellum has a layered geometry and consists of four distinct layers – the oligodendrocyte, granular, Purkinje–Bergmann (PB) and molecular layers – as identified in previous studies [16, 20] (Figure 2A). To investigate transcription factor (TF)-target and ligand-receptor gene interactions, we selected 39 TFs and 773 target genes and 23 ligand genes and 26 receptor genes, respectively, from the GASTON spatially variable genes [16].

We used GLACIER to focus on two biological questions, which were previously out of reach for existing methods studying cellular interactions:

1. What is the *directionality* of cellular interactions – and more generally, information flow – within a layer?
2. Is there a *discontinuity* in information flow across layers?

To answer the first question, we used GLACIER to identify cellular interactions *within* each cerebellum layer (e.g. granule layer, Figure 2B), which involves forming a spatial DAG (Figure 2C) oriented according to the GASTON isodepth *d* which smoothly varies within each layer. Importantly, by *inverting* the isodepth (Methods, Figure 2D), GLACIER identifies cellular interactions in the opposite direction and thus determine the *directionality* of cellular interactions. For the second question, we used GLACIER to identify cellular interactions that are unique to cells at the *boundary* of a cerebellum layer (Figure 2E).

**Figure 2:**
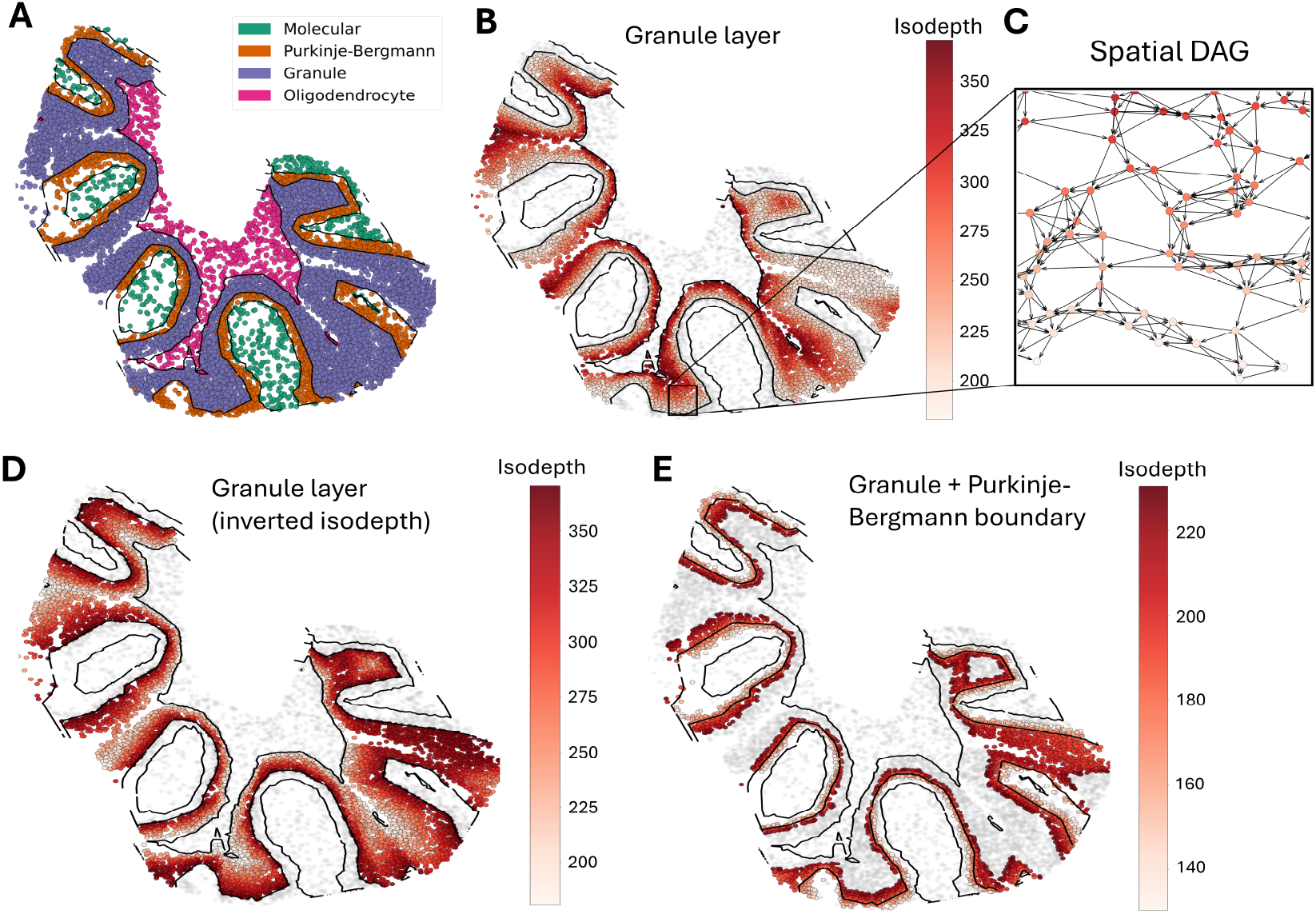
**(A)** Slide-SeqV2 mouse cerebellum dataset with each cell colored by its layer assigned by GAS-TON [16]. Each layer is named according to its dominant cell type as in [20, 35]. **(B)** Cells in granular layer colored by GASTON isodepth. Black curves denote GASTON-idenfied layer boundaries. **(C)** Spatial DAG *G* = (*V, E*) constructed by GLACIER in granular layer, with directed edges (*i, j*) oriented from high isodepth to low isodepth, i.e. edge *i j* →if *d*_*i*_ *> d* _*j*_. **(D)** Inverted isodepth 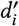 (equation (6)) in granular layer, which allows for identification of *bidirectional* cellular interactions by GLACIER. **(E)** Cells with isodepth *d*_*i*_ within 50*μm* of layer boundary between granular layer and Purkinje-Bergmann layer, colored by isodepth *d*_*i*_, allowing for GLACIER to identify cellular interactions *between* cell types.

To address these questions, we analyzed the data at multiple levels. We identified frequently-regulated target genes and performed gene set enrichment analysis (GSEA) to determine which biological processes were enriched among these targets [36]. We further examined specific TF-target and ligand-receptor gene pairs to explore key regulatory interactions at fine resolution. Lastly, we also exploited GLACIER-derived lag estimates to identify ligands with short- and long-ranging activity at layer boundaries.

### 3.1 Gene sets

#### Direction of information flow in oligodendrocytes

We examined gene set enrichments for oligodendrocytes in both orientations of isodepth. We observed that increasing isodepth, i.e. where the DAG leads away from the granular layer, exhibited a much more coherent and interpretable set of enriched biological processes. This direction was strongly associated with axonogenesis, synaptic transmission, and neuronal connectivity, suggesting that oligodendrocytes in this orientation actively shape neuronal function rather than simply responding to their environment (Figure 3A-B). Function enrichments included axon development, regulation of neurotransmitter secretion, and neuron projection morphogenesis, all of which align with known roles of oligodendrocytes in supporting and refining neural circuits [37].

**Figure 3:**
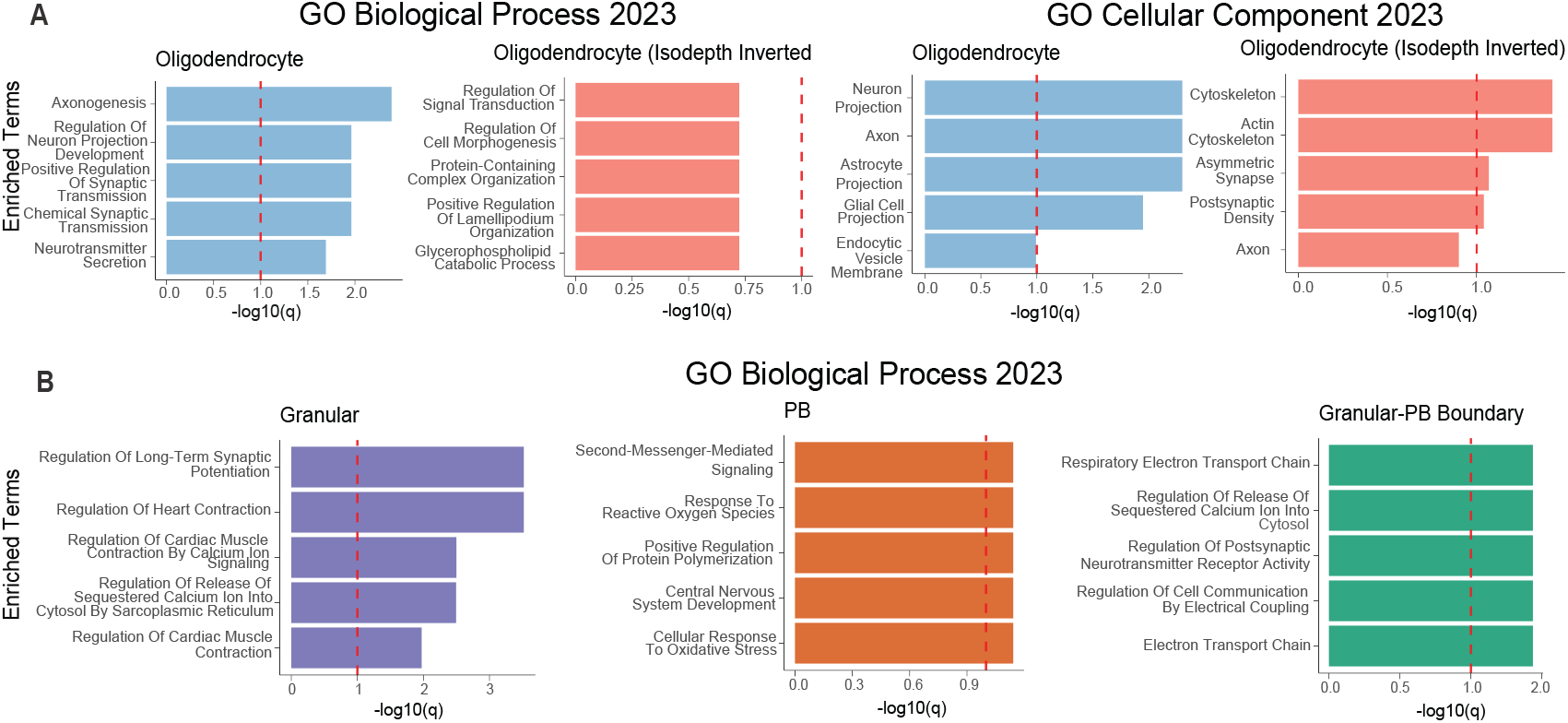
**(A)** Gene set enrichment analysis (GSEA) with Enrichr [36] for cellular interactions identified by GLACIER in the oligodendrocyte layer with isodepth *d* and inverted isodepth *d*^’^ isodepth orientations for biological process (left) and cellular component (right) gene sets. **(B)** GSEA for cellular interactions identified by GLACIER in the granular layer (left), Purkinje-Bergmann (PB) layer (middle), and the boundary between the granular and PB layers (right).

At the gene level, this enrichment was driven by regulators of axon guidance and synaptic adhesion, such as neurexin-1 (*Nrxn1*) and neurofascin (*Nfasc*), which mediate neuron-glia communication, and *Mag*, which plays a key role in stabilizing axonal connections. In contrast, the decreasing isodepth (inverted) direction showed weaker and less functionally coherent enrichments, with a shift toward intracellular and cytoskeletal remodeling rather than neuron-facing processes. Genes such as *Cfl1* and *Phip*, which regulate actin dynamics [38] and intracellular signaling, were more prominent in this direction, suggesting a greater emphasis on structural maintenance or adaptation.

These findings raise the possibility that oligodendrocytes’ instructive role proceeds in the direction of isodepth, with less regulatory control exerted in the other direction. Further experimental work would be needed to investigate this hypothesis. Indeed, the value of GLACIER is precisely in helping generate such hypotheses for further investigation.

#### Regulatory discontinuity at the boundary of granular and Purkinje-Bergmann layers

Gene set enrichment at this boundary revealed a distinct functional signature that differs from either layer alone. While the granular layer was enriched for processes related to long-term synaptic potentiation and calcium-mediated excitability (Figure 3C, left), and the PB layer showed enrichments for oxidative stress response and second-messenger signaling (Figure 3C, middle), the boundary exhibited a distinct focus on synaptic modulation, electrical coupling, and metabolic activity (Figure 3C, right). This suggests that cells at the interface of the granular and Purkinje-Bergmann layers play a specialized role in mediating communication and bioenergetic demands between the two layers.

One of the strongest enrichments at the boundary included regulation of postsynaptic neurotransmitter receptor activity and cell communication by electrical coupling, suggesting rapid synaptic transmission. We also observed enrichment of electron transport chain activity possibly due to metabolic interactions. The gene set included *Dlgap1* and *Nptx1*, both of which are involved in synaptic scaffolding and activity-dependent synapse formation, reinforcing the idea that the boundary plays a role in synaptic refinement and communication [39]. In addition, *Ndufa5* and *Sdha*, key components of the mitochondrial respiratory chain [40], indicate that cells in this region experience high metabolic demand and are likely to support sustained neurotransmission.

These results suggest that the Purkinje-Bergmann and granular cell boundary is not simply a gradient between two transcriptional programs, but rather a specialized interface with its own regulatory demands. Experimental validation could determine whether these transcriptional differences are critical to sustaining high-frequency signaling and synaptic communication in the cerebellum.

#### Gene pairs

GLACIER reveals several biologically meaningful pairs of genes (TF-target or ligand-receptor) with spatial Granger causal relationships within individual layers. For example, GLACIER identifies that the TF *Hspa5* is spatially Granger causal for the target gene *Pink1* (Figure 4A), with *Pink1* expression closely following *Hspa5* expression. *Hspa5* maintains endoplasmic reticulum (ER) homeostasis [41] while *Pink1* performs mitochondrial quality control [42], suggesting that activation of general ER regulation results in recruitment of proteins that regulate specific cellular functions. GLACIER also identifies that the ligand *S100b* is spatially Granger causal of the receptor *Ptprz1* (Figure 4B). *S100b* has been suggested to support neuron migration [43] while *Ptprz1* has been observed to modulate cell migration in some tissues [44], suggesting that neurons in the granular layer may be moving in the isodepth direction towards the adjacent oligodendrocyte layer.

**Figure 4:**
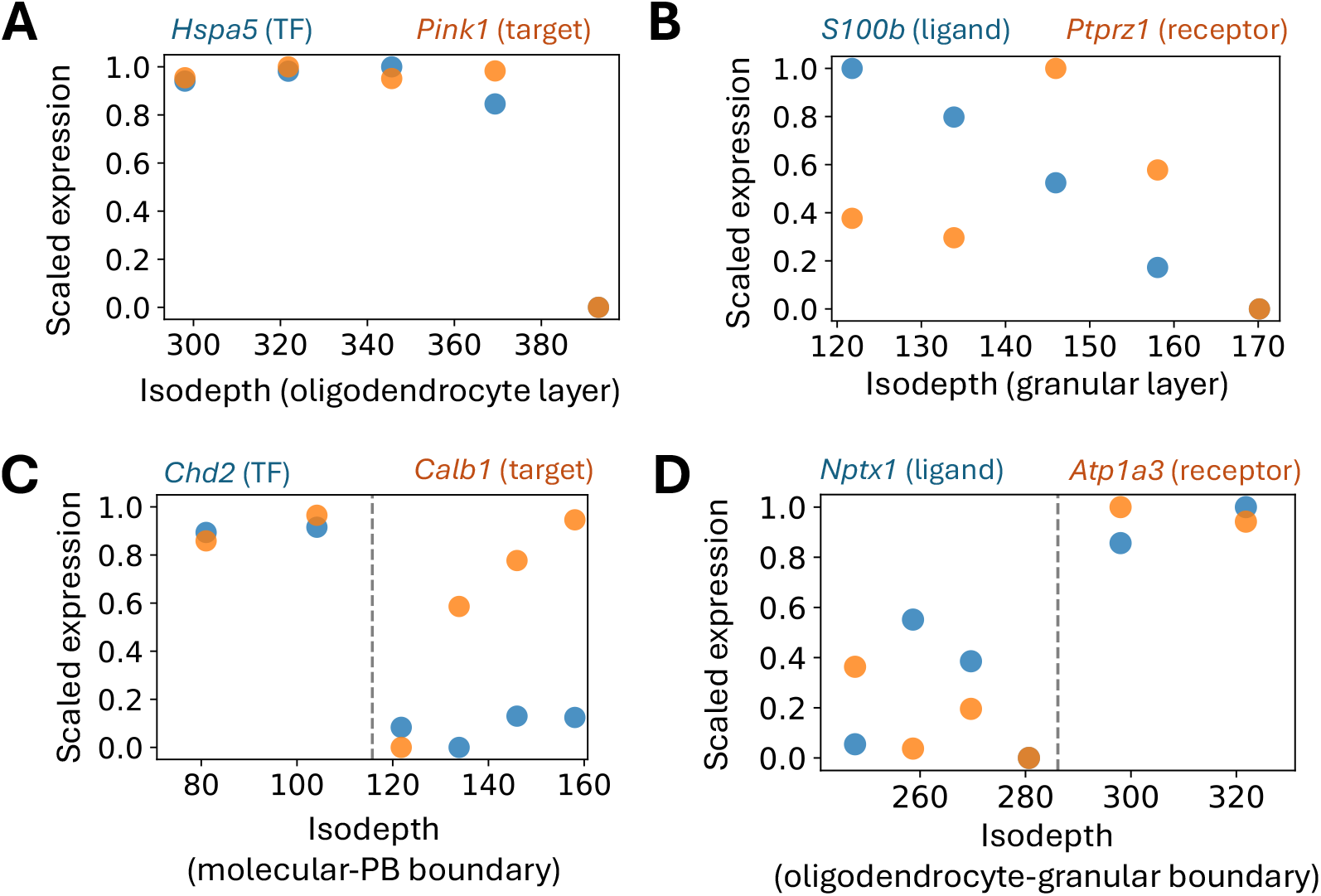
Isodepth versus scaled expression for **(A)** transcription factor *Hspa5* and target gene *Pink1* in the oligodendrocyte layer, **(B)** ligand *S100b* and target gene *Ptprz1* in the granular layer, **(C)** transcription factor *Chd2* and target gene *Calb1* at the boundary between the molecular and Purkinje-Bergmann (PB) layers, and **(D)** ligand *Nptx1* and receptor *Atp1a3* at the boundary between the oligodendrocyte and granular layers.

Further, GLACIER identifies spatially Granger causal pairs of genes that are unique to the *boundary* between two layers. For example, GLACIER identifies an interaction between the TF *Chd2* and target gene *Calb1* at the boundary between the molecular and Purkinje-Bergmann (PB) layers (Figure 4C). *Chd2* has been observed to broadly regulate neuronal function [45] while *Calb1* is a marker gene for Purkinje cells [46], suggesting that *Chd2* may be regulating the function of Purkinje cells specifically at the molecular-PB layer boundary. GLACIER also identifies an interaction between the ligand *Nptx1* and the receptor *Atp1a3* that uniquely occurs at the boundary of the oligodendrocyte and granular layers; both *Nptx1* and *Atp1a3* are known to be involved in neuronal signaling and excitation [47, 48], suggesting potential communication between oligodendrocytes and granular cells at the boundary.

In this way, GLACIER identifies cellular interactions within and across layer boundaries which cannot be found by existing approaches which do not incorporate the global tissue topography.

#### Spatial range estimation via GLACIER lag estimates

GLACIER’s causal inference framework reports not only the regulatory interactions but also the lag at which the interaction operates (Methods). This allows us to estimate the spatial range of a regulator gene (i.e. a TF or ligand), with bigger lags corresponding to a larger spatial range of a regulator. We investigated some settings where GLACIER reported bimodal lags across TF or ligand regulators (Figure S1), denoting substantial variations in the spatial ranges of regulators.

##### Molecular Layer/Purkinje-Bergmann Boundary

The boundary between the Molecular and Purkinje-Bergmann layers displayed particularly pronounced bimodality in the ligand distance distribution. The longest-ranging ligand at this interface was *Apoe*, a gene well-known for its role in lipid transport, synaptic plasticity, and neuronal maintenance [49] *Apoe*’s ability to bind lipoproteins and facilitate cholesterol uptake is crucial for neuronal repair and may underlie its long-range effects across these layers. Conversely, the shortest-acting ligand we identified was *Gdf10*, a member of the TGF-*β* superfamily involved in neuronal regeneration and repair [50]. *Gdf10*’s localized expression and role in modulating cell growth suggest its influence is confined to more proximal targets in neighboring cells.

##### Purkinje-Bergmann/Granular Layer Boundary

We also observed substantial spatial-range variation between ligands at the Purkinje-Bergmann/Granular layer boundary. Here, the furthest-acting ligand was *Gng13*, a *G*-protein gamma subunit that partners with alpha and beta subunits to regulate a variety of downstream signaling pathways [51]. Its presence in long-range signaling contexts suggests that *Gng13* may orchestrate interactions spanning multiple cell types or spatial zones in the cerebellar architecture. While there was no single dominant short-range ligand at this boundary, the *Thy1* receptor was repeatedly seen. *Thy1* (also known as *Cd90*) is a cell-surface glycoprotein involved in cell adhesion, neurite outgrowth, and T-cell activation [52]. Its role in modulating cell-cell contact may explain its consistent appearance among ligands operating over limited spatial distances, hinting that *Thy1*-dependent interactions mediate localized communication in these cerebellar layers.

Taken together, these findings underscore how spatial organization can drive bimodal patterns of ligand-receptor signaling: some factors, like *Apoe* and *Gng13*, can reach distant targets, while others, such as *Gdf10, Gnas*, and *Thy1*, function in tight, localized niches. By systematically exploring these “lag distributions,” GLACIER reveals how cerebellar architecture shapes the breadth and specificity of cellular communication.

## 4 Discussion

GLACIER introduces a framework for inferring causal transcriptional and signaling relationships in spatial transcriptomics by leveraging a global tissue topology. Unlike previous approaches that rely on local cellcell adjacency, GLACIER constructs a directed acyclic graph (DAG) based on a global spatial coordinate, allowing us to infer how regulatory programs unfold along natural tissue axes. By orienting regulatory interactions along inferred spatial axes, we move beyond local spatial correlations and instead model how gene regulation and cell signaling propagate through *global* tissue structure. Applied to the cerebellum, GLACIER uncovered directional transcriptional programs within layers and distinct regulatory activity at layer boundaries, demonstrating its ability to identify spatially organized cellular interactions.

One limitation of GLACIER is that it requires tissues where a global spatial coordinate, such as isodepth, can be inferred. However, this is not a fundamental restriction – user-guided segmentation of complex tissues can enable locally defined spatial coordinates, allowing GLACIER to be applied more broadly. Moreover, one can also use a supervised approach for determining such spatial coordinates, such as in [53, 54]. Another challenge is transcript sparsity, a common issue in spatial transcriptomics that may become more pronounced with technologies like Xenium [55]. However, GLACIER naturally mitigates this issue by aggregating over multiple cells (i.e. cells with same isodepth), increasing robustness to sparsity. Additionally, GLACIER’s bootstrap-based hyperparameter selection provides a systematic way to ensure robust inferences.

While we applied GLACIER’s spatial Granger causality approach to brain layers, the methodology generalizes to other structured tissues. Future work could leverage this framework to study how signaling cascades change radially from blood vessels or how the interior and periphery of a tumor microenvironment exhibit distinct regulatory states. Looking forward, integrating GLACIER with spatial proteomics [56] and multimodal single-cell datasets [57] could provide a more comprehensive view of spatially structured regulatory programs, further enhancing our ability to decode the organization of complex tissues.

## Acknowledgments

U.C. was supported in part by funding from the Eric and Wendy Schmidt Center at the Broad Institute of MIT and Harvard. H.L. and R.S. were partially supported by the Chan Zuckerberg Initiative.

## Appendix

## A Robustness testing

Velorama sweeps over a range of regularization hyperparameters (*λ*), averaging TF-target gene interactions from models where the percentage of non-zero interactions fall within a given threshold (by default, 0.01 to 0.95). To find *λ* values meeting this threshold for each layer and isodepth orientation, we increased the number of *λ*s tested from the default of 10 to 30. We also narrowed the search range from (0.01, 10) to (0.001, 0.1), since we found suitable *λ*s was always below 0.1 empirically. We used a higher learning rate (*η* = 0.1) for TF-target gene analysis and a lower learning rate (*η* = 0.01) for ligand-receptor analysis. The changes in learning rate were made across all runs, and the number of iterations per *λ* during training was significantly increased from the default of 1000 to between 10,000 and 30,000, depending on the run.

We verified other hyperparameters by testing on stratified thirds of the dataset. Each stratified third was constructed by evenly selecting cells from cell type-specific subsets and then combining them, ensuring that the cell type distribution was preserved. We tuned hyperparameters so that results were consistent across stratified thirds and the overall dataset (for both normal and inverted isodepth). Finally, we analyzed the loss for each run to confirm smooth training behavior and convergence at the end of each model training run.

**Figure S1:**
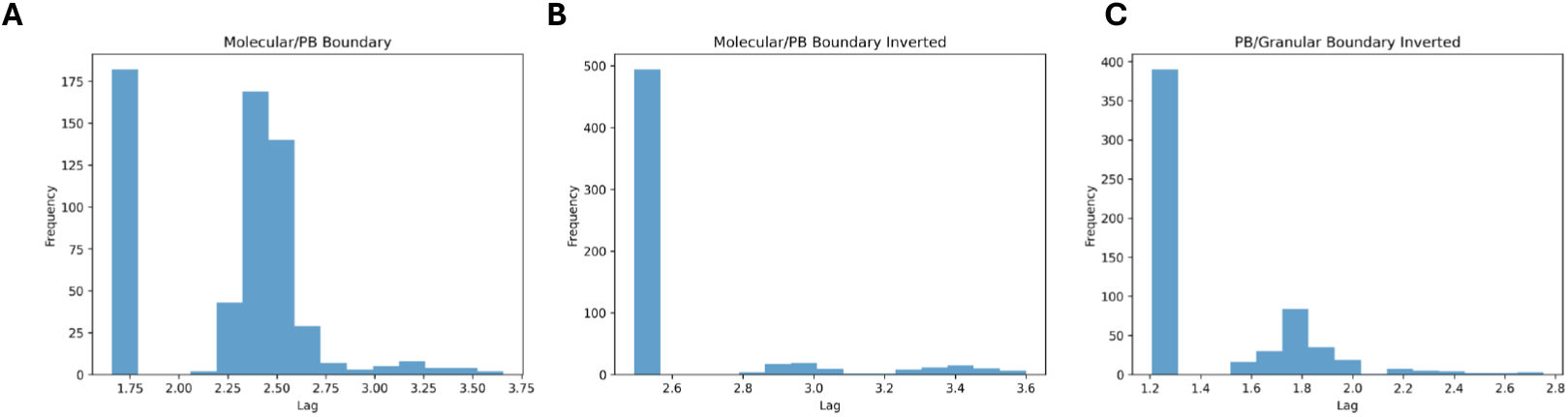
Distribution of lags identified by GLACIER across all candidate interactions in **(A)** the boundary between the molecular and Purkinje-Bergmann (PB) layers using the isodepth *d*, **(B)** the boundary between the molecular and PB layers using the inverted isodepth *d*^’^, and **(C)** the boundary between the PB and granular layers using the inverted isodepth *d*^’^.

